# Metacognitive Efficiency Reduces Confirmation Bias in Perceptual Decision Making

**DOI:** 10.64898/2026.06.18.733181

**Authors:** Alexis Pérez-Bellido, Rubén Moreno-Bote, Lluís Fuentemilla

## Abstract

Humans exhibit a pervasive drive toward self-consistency, often failing to revise previous decisions even when confronted with contradictory evidence. Here, we investigate the computational mechanisms underlying decision revision in perceptual tasks, examining the regulatory role of metacognition. To do so, we capitalize on a novel paradigm in which participants are repeatedly presented with identical sensory information and allowed to revise their choices after each exposure. Our results reveal that repeated exposure to the same stimulus systematically biases subsequent judgments toward prior responses. Using drift-diffusion modeling, we tested competing explanations incorporating different assumptions about how prior choices affect evidence accumulation. Our findings indicate that consistency biases emerge from asymmetric sensory weighting, selectively amplifying information consistent with previous choices—a phenomenon akin to confirmation bias. Crucially, individuals with higher metacognitive skills exhibited weaker confirmatory biases and more flexible integration of repeated sensory information, enabling greater adaptability in decision-making. These findings highlight the continuous nature of perceptual inference and underscore metacognition’s pivotal role in mitigating bias and optimizing decision flexibility.

## Introduction

“Humans are the only animals that trip twice over the same stone.” While perhaps scientifically hyperbolic and inaccurate, this popular Spanish proverb encapsulates a central paradox of human cognition: our persistent inclination to adhere to prior beliefs and choices even when faced with new, contradictory information. A key challenge in decision science is understanding why humans often fail to adequately update their judgments based on evolving evidence. For instance, after forming an initial opinion about a film, does rewatching it typically lead to a revision of that judgment—or are first impressions inherently resistant to change? Empirical evidence indicates that initial judgments, once formed, are remarkably stable and challenging to overturn (Rabin & Schrag, 1999; Stone et al., 2022). Studies in perceptual decision-making consistently demonstrate that humans favor interpretations that align with previous subjective decisions, exhibiting systematic biases toward initial categorical judgments. These biases have been modeled as the result of a self-consistent “Bayesian” observer (Luu & Stocker, 2018, 2021), wherein biased decisions emerge from a top-down inferential process; in this framework, the readout of low-level sensory estimates stored in short-term memory is conditioned by earlier high-level judgments. A parallel line of research, employing intermittent decision-making paradigms—where individuals sequentially process sensory information and make preliminary categorical judgments midway—has provided an alternative explanation for this self-consistency bias. These studies propose that biases may emerge from the asymmetric encoding of sensory inputs, a phenomenon mediated by a selective-gain mechanism through which participants selectively increase sensitivity to evidence aligned with prior decisions (Bronfman et al., 2015; Kaanders et al., 2022; Rollwage et al., 2020; Talluri et al., 2018, Talluri et al., 2021).

Collectively, these findings underscore a human proclivity toward consistency that is evident even in tasks where perceptual evidence is uncorrelated across trials. Such biases extend beyond perceptual tasks to influence short-term memory and decision-making, as demonstrated by serial dependency phenomena (Fiser et al., 2016; Fritsche et al., 2017; Abrahamyan et al., 2016; Urai et al., 2019). For example, Urai et al. (2019) showed that participant choices were biased toward previous decisions even when trials were independent—a tendency captured by drift-diffusion model parameters reflecting confirmation-like processes. This general drive for self-consistency constitutes a hallmark of confirmation bias. Confirmation bias is the cognitive tendency to search for, interpret, favor, and recall information that confirms one’s pre-existing beliefs, while ignoring or discounting contradictory evidence, (Nickerson, n.d.). This cognitive heuristic has profound negative consequences at a societal level, as illustrated by the contemporary spread of fake news and disinformation (Lazer et al., 2018). Why, then, is human cognition structured around such a seemingly maladaptive bias? From a bounded rationality perspective, evidence suggests that at the individual level, confirmatory biases may confer adaptive advantages. Reinforcement learning (RL) models incorporating confirmation biases have been shown to outperform unbiased algorithms, suggesting these biases efficiently disregard uninformative, stochastic negative prediction errors and optimize resource accumulation (Lefebvre et al., 2022; Palminteri & Lebreton, 2022; Tarantola et al., 2021). Furthermore, modeling work using the “self-consistent” Bayesian observer hypothesis has demonstrated that, if an initial decision is correct, being self-consistent confers cognitive advantages by preserving stable representations of the environment and protecting working memory contents from corruption by subsequent noise (Qiu et al., 2020).

Thus, to take advantage of a self-consistent strategy, correctly identifying when one is correct or incorrect becomes crucial. This ability to evaluate one’s decision accuracy is known as metacognition (Fleming, 2017; Fleming & Dolan, 2012; Fleming & Lau, 2014). In perceptual decision-making tasks, metacognition is typically measured by the extent to which the strength of one’s confidence is coupled with objective performance. The impact of confidence on the perceptual decision process is well-documented; it acts as an internal control signal, influencing how past information guides current behavior and regulating how new information is integrated (Balsdon & Philiastides, 2024; Peters et al., 2017; Rollwage et al., 2020). Research indicates that decisions made with high confidence are particularly resistant to revision. Indeed, when a decision-maker is highly confident, the brain amplifies the processing of new evidence that confirms the initial choice while effectively abolishing the processing of disconfirmatory evidence (Balsdon et al., 2020; Desender et al., 2018; Rollwage et al., 2020).

Nevertheless, while theoretical models suggest that individuals with higher metacognitive abilities might better harness the stabilizing benefits of confirmation bias (Rollwage et al., 2020), empirical evidence exploring the interaction between metacognition and confirmation bias remains limited.

In the present study, we investigated the role of metacognitive efficiency as a potential mitigating factor against confirmation bias in decision-making. To test this hypothesis, we developed a novel experimental paradigm. Specifically, participants encountered a sequence of oriented gratings and were required to judge whether the sequence contained, on average, more cardinal or diagonal orientations. Following their initial decision, they were repeatedly presented with the identical stimulus sequence for two to three additional repetitions, providing the opportunity to revise their judgment up to two times per trial.

This design aimed to simulate a particular real scenario, which is rarely experimentally tested, where individuals are exposed to the same information on multiple occasions. Crucially, this approach allowed us to distinguish our findings from existing models: unlike paradigms that examine how categorical choices bias the retrieval of information from short-term memory (Luu & Stocker, 2018, 2021) or how ongoing decisions are interrupted midstream (Bronfman et al., 2015; Talluri et al., 2018), our design focused on the stability of judgments across repeated exposures to the same sensory evidence.

Using this framework, we first examined whether the tendency to repeat a response stemmed from a simple repetition bias (a non-sensory preference for consistency) or was instead characterized by an asymmetric weighting of sensory evidence, where confirmatory information is prioritized over disconfirmatory input. Second, we collected confidence judgments after each decision. By pairing these subjective reports with objective performance, we calculated participant-level metacognitive efficiency (Fleming, 2017; Maniscalco & Lau, 2012). We specifically predicted that higher metacognitive efficiency would reduce the probability of exhibiting confirmation biases. Furthermore, we hypothesized that the strength of the confirmation bias itself was modulated by individual differences in metacognitive ability (Rollwage & Fleming, 2021). Consequently, we expected that participants with higher metacognitive capacities would demonstrate superior information integration across repetitions, resulting in more substantial improvements in overall task performance.

## Results

In Experiment 1, our primary objective was to characterize the computational nature of decision biases—specifically, to determine whether the tendency to repeat a choice stemmed from a non-sensory repetition bias or an asymmetric weighting of sensory evidence. To this end, we employed a perceptual decision-making task involving oriented Gabor gratings (Fig. 1A). Participants were presented with a sequence of six stimuli and were required to categorize the sequence as containing, on average, a majority of orientations closer to the cardinal or the diagonal axis. A defining feature of this paradigm was the use of identical repeated sequences: following the initial categorization of a sequence (P1), the exact same physical stimulus sequence was repeated up to three times. Participants were explicitly informed that the sequences were identical and were encouraged to use these subsequent presentations to either confirm or revise their prior judgments. Performance feedback was provided only at the conclusion of each trial. Task difficulty was parameterized by a Decision Variable (DV), where positive values indicated a diagonal majority and negative values indicated a cardinal majority; values near zero represented high sensory ambiguity. To maintain stable performance across participants, a 1-up/2-down staircase procedure was employed to modulate the DV. Crucially, to prevent choice-induced changes in difficulty, the staircase was updated based exclusively on the accuracy of the response to the first presentation (P1). Experiment 2 aimed to assess how individual metacognitive abilities modulated the tendency to repeat choices. While the core paradigm remained the same as in Experiment 1, participants in Experiment 2 also reported their subjective confidence following each categorical judgment using a Likert scale. To maximize the trial count for robust metacognitive estimation, trial sequences in this experiment were presented only twice (P1 and P2), allowing participants to change their mind once per trial. These confidence reports, paired with objective accuracy, enabled the calculation of individual metacognitive efficiency.

**Figure 1.**
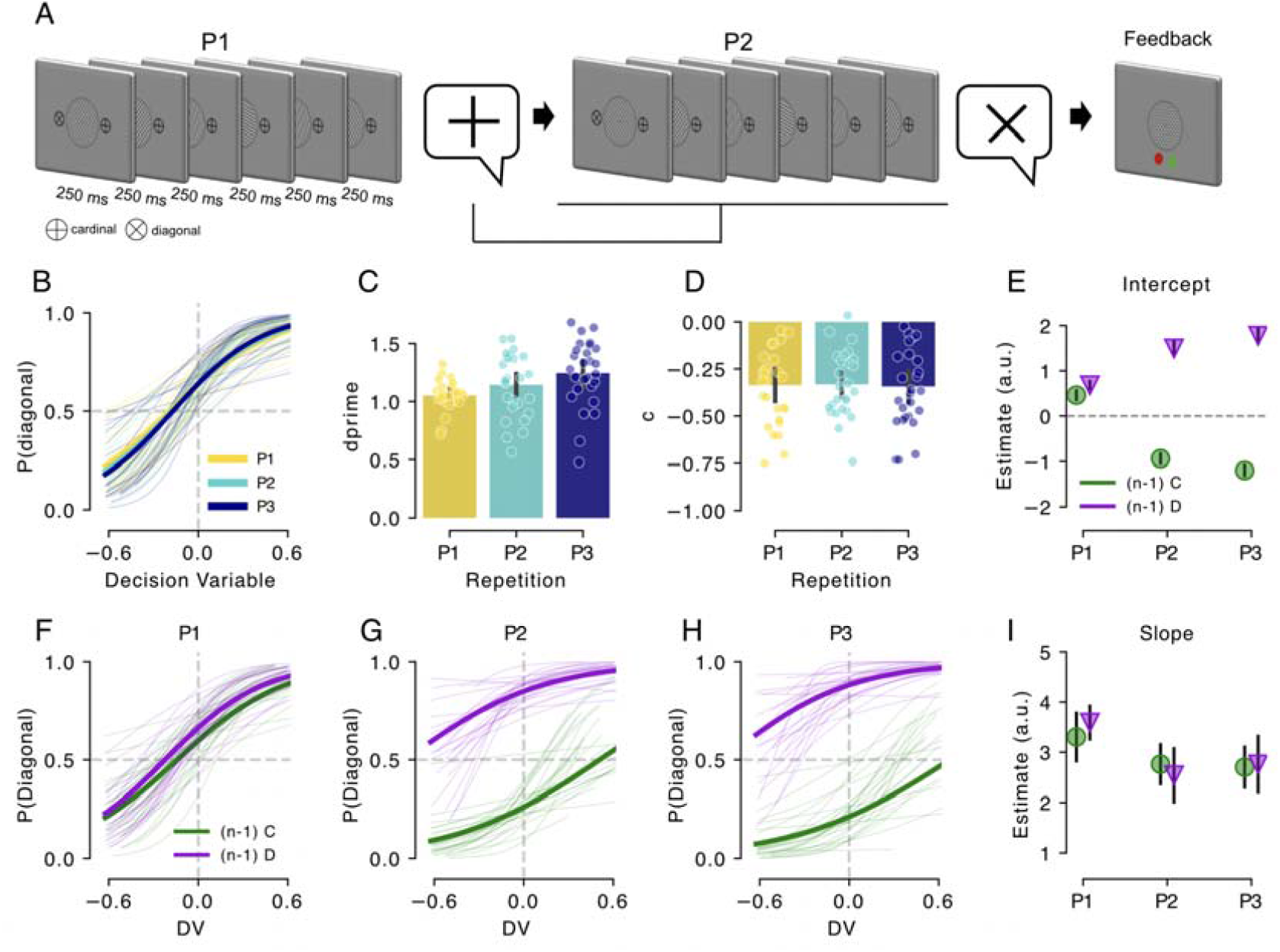
Behavioral performance and choice-conditioned psychometric shifts across identical repetitions. A, Experimental paradigm. Schematic of a single trial with two presentations (note that Experiment 1 utilized three presentations) showing the sequential presentation of identical oriented gratings (P1 and P2). Following each sequence, participants categorized the predominant orientation as cardinal or diagonal. Response mappings (icons) were displayed concurrently and randomized across presentations to decouple sensory judgments from motor preparation. Feedback was provided at the end of each trial. B, Psychometric curves representing the probability of a “diagonal” categorization as a function of the Decision Variable (DV) for the three sequential stimulus presentations (P1, light yellow; P2, cyan; P3, dark blue). Thin lines represent individual participants; thick lines represent the group-level logistic fit. C–D, Signal detection theory (SDT) metrics across repetitions. Sensitivity (d’; c) shows a significant increase from P1 to P2 and P3, reflecting information accumulation, while the decision criterion (c; D) illustrates a persistent negative bias toward the cardinal category. Points represent individual subjects; error bars represent ±1 s.e.m. E, Intercept estimates from the hierarchical mixed-effects model, quantifying systematic bias as a function of choice history (n-1). Estimates are conditioned on whether the previous response was cardinal (circles) or diagonal (triangles). Error bars represent 95% confidence intervals. F–H, Choice-conditioned psychometric curves for the first (F), second (G), and third (H) presentations, split by the preceding choice (n-1; cardinal, orange; diagonal, blue). The horizontal shift captures the influence of the previous judgment on current perception. I, Slope estimates from the hierarchical mixed-effects model, representing sensitivity to the sensory DV across repetitions, conditioned on the previous response. The decrease in slope magnitude from P1 to P3 corresponds to a choice-conditioned loss in precision that is not captured by d’ metrics. Together, these panels demonstrate that while redundant exposure improves absolute sensory precision (d’), it simultaneously entrains a robust consistency bias and a specific loss in sensitivity to sensory evidence that contradicts the previous judgment.

### Participants integrate information across repetitions

Participants’ ability to discriminate stimulus orientation improved with repeated exposure. This was evidenced by the steeper slope of the psychometric curve at the third presentation (P3) compared to the first (P1) and second (P2), as shown in Fig. 1A. Signal detection theory analyses supported this pattern, with d’ values increasing across repetitions (Fig. 1C; F(2, 52) = 15.87, p < 0.001, ηG^2^ = 0.10). Criterion analyses revealed a systematic bias toward categorizing stimuli as diagonal rather than cardinal, suggesting that participants required strongly horizontal or vertical orientations to judge the mean orientation as cardinal. Notably, this bias remained stable across presentations Fig. 1B; F(2, 52) = 0.17, p = 0.843, ηp^2^ < 0.001). To examine the dynamics of response bias more closely, we fitted a generalized linear mixed model (GLMM) to determine whether participants’ current responses were influenced by their previous choices (Fig. 1F-H). Changes in the intercepts of the psychometric curves revealed that participants’ responses in P2 and P3 were biased toward their immediately preceding choices (Fig. 1E). For instance, if a participant categorized the sequence as cardinal in P1, they were more likely to repeat that categorization in P2. Importantly, no such carryover effect was observed between different trials: participants’ P1 responses were not influenced by their P3 decisions from the previous trial. These results indicate that decision-related information is actively integrated within— but not across—trials, with participants combining prior choices and current perceptual evidence to improve their judgments. We also found out that when psychometric curves were separated based on prior choices, the slopes for P2 and P3 were significantly shallower than for P1, as evidenced by the change in the DV slope (Fig. 1I). This suggests that although repeated exposure generally led to better performance, the influence of prior decisions reduced participants’ sensitivity to new sensory evidence in later presentations.

### Biased Integration of Perceptual Evidence Under Repeated Exposure

We next sought to understand how participants integrated information across repetitions. Several non-mutually exclusive hypotheses could account for the observed effects. One possibility is that participants simply repeated their choices during the second and third presentations when they were highly confident in their initial decision. Such a strategy would not necessarily alter how new information is integrated during subsequent repetitions. Alternatively, participants might exhibit a bias in the integration process itself—specifically, a systematic asymmetric accumulation of evidence favoring previous choices (Rollwage et al., 2020; Urai et al., 2019). These hypotheses cannot be disentangled on the basis of accuracy alone, as both can produce similar patterns of response bias. However, reaction times (RTs) provide additional insight into the underlying integration process. Response repetition biases typically affect only fast responses, whereas integration biases influence both fast and slow responses. To distinguish between these mechanisms, we employed Drift Diffusion Models (DDMs), which jointly model choice accuracy and RTs (Ratcliff et al., 2016; Ratcliff & McKoon, 2008).

In DDMs, perceptual decisions arise from the noisy accumulation of sensory evidence over time until a decision threshold is reached. Key parameters include the drift rate (v), reflecting the speed of evidence accumulation; the decision threshold (a), indicating the amount of evidence required for a response; the starting point (z), representing an initial bias prior to evidence accumulation; and the non-decision time (Ter), capturing sensory and motor delays. A tendency to repeat previous choices would manifest as a shift in the starting point, reflecting a bias present before evidence accumulation begins. In contrast, an integration bias—where prior choices modulate the processing of new evidence—would be reflected in changes to the drift rate, for example via a shift in the drift criterion that biases accumulation toward the boundary corresponding to the favored choice. We fitted hierarchical DDMs (Wiecki et al., 2013) to participants’ data across repetitions and compared models incorporating different bias mechanisms. The best-fitting model indicated that drift rate varied as a function of choice history, suggesting that participants’ integration of new evidence was biased toward their previous choices (Fig. 2A). Indeed, DDM parameter estimates showed that the drift criterion parameter was increasingly biased toward the previous choice in each presentation, indicating that the integration of evidence into a decision was conditioned on participants’ prior choices (Fig. 2B and C). However, these results did not clarify whether the biased drift rate reflected an information-independent shift in accumulation toward one response boundary or an asymmetric weighting of sensory samples depending on their consistency with the previous choice (Bronfman et al., 2015; Peters et al., 2017; Rollwage et al., 2020; Talluri et al., 2018). To adjudicate between these explanations, we conducted simulations in which sensory samples were processed through a psychometric kernel prior to accumulation. To simulate an information-independent drift rate bias, we shifted the observer’s psychometric function toward the previous choice on the second presentation (Fig. 2D), thereby biasing sample integration monotonically toward that choice regardless of sample content. To simulate asymmetric integration, we instead implemented a lapse mechanism selectively affecting samples inconsistent with the previous choice (Fig. 2F). This model emulates an integrator with reduced sensitivity to disconfirmatory evidence (or, equivalently, increased sensitivity to confirmatory evidence).

**Figure 2.**
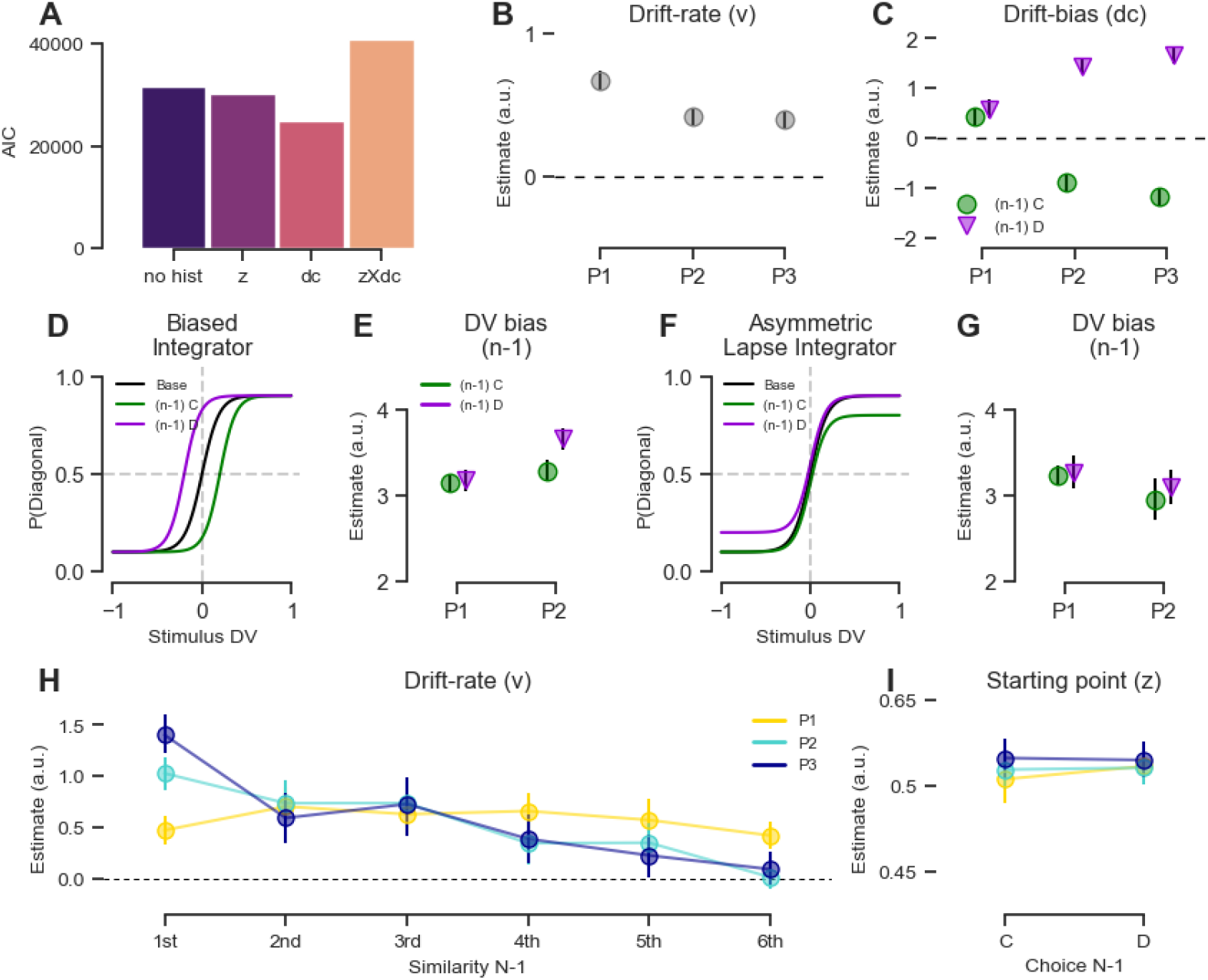
Computational mechanisms of confirmation bias under repeated sensory exposure. A, Model comparison using the Akaike Information Criterion (AIC). Hierarchical Drift-Diffusion Models (HDDMs) were fitted to adjudicate between distinct mechanisms of choice-history integration. The model incorporating history-dependent shifts in the drift criterion (dc) provided the best fit, outperforming models assuming no history dependence (“no hist”), a history-dependent starting point (z), or a combination of both (zXdc). B, Estimated mean drift rate (v) across three sequential presentations (P1, P2, P3). The baseline rate of evidence accumulation decreases following the initial exposure. C, Drift bias (dc) parameter estimates conditioned on the observer’s previous choice (n-1): Cardinal (C, green circles) or Diagonal (D, purple triangles). The accumulation process becomes increasingly biased toward the boundary of the preceding choice across identical repetitions (P2 and P3). D–G, Qualitative comparison of integration mechanisms via simulation. Panels D and E illustrate a biased integrator where choice history induces a horizontal threshold shift; note that choice-dependent sensory sensitivity (slope) increases across repetitions. Panels F and G illustrate an asymmetric lapse integrator, where choice-dependent integration failures effectively reduce sensory precision (DV bias), reproducing the empirical sensitivity loss observed in Fig. 1I. H, Drift rate (v) as a function of sensory sample similarity to the previous choice, ranked from most similar (1st) to least similar (6th). During repeated exposures (P1, yellow; P2, cyan; P3, dark blue), participants exhibit asymmetric weighting: samples consistent with the initial choice are weighted heavily, whereas inconsistent (disconfirmatory) samples are significantly underweighted. I, Starting point (z) estimates conditioned on the previous choice. The starting point remains near the unbiased baseline (0.5), confirming that the observed self-consistency effect is driven by biased evidence integration rather than a pre-accumulation response bias. Error bars represent 95% confidence intervals.

Both simulated mechanisms enhanced performance in the second presentation (P2) and reproduced a bias to repeat the previous choice (Fig. S2 and S3). However, they made distinct predictions regarding the slope of choice-dependent psychometric functions. The biased integrator produced steeper slopes in P2 relative to P1, reflecting a global shift in accumulation (Fig. 2E). In contrast, the lapse integrator produced shallower slopes in P2 relative to P1 (Fig. 2G). Crucially, only the lapse integrator replicated a key feature of the empirical data: the slope of the choice-dependent psychometric function was reduced in P2 compared to P1 (Fig 1I). These results indicate that evidence accumulation is not simply shifted toward a preferred boundary; rather, observers appear to underweight sensory samples that contradict their initial choice, resulting in reduced sensitivity to repeated evidence.

### Confirmation Bias results from asymmetric evidence weighting

We hypothesized that this suboptimal sampling pattern reflects a confirmation bias in perceptual integration. To test this hypothesis, we fitted a hierarchical multiple-regression DDM to determine whether the similarity between individual sensory samples and the previous choice predicted their contribution to the drift rate. Within each presentation, stimuli were reordered according to their distance in Decision Variable (DV) space from the previous choice. The first regressor contained the DV values of samples most similar to the previous choice, while subsequent regressors represented samples with increasing distance, such that the final regressor included the most dissimilar (disconfirmatory) samples. As a baseline, we fitted an equivalent model for P1 using the participant’s response from the final presentation (P3) of the preceding trial. Consistent with confirmation-biased integration, samples more similar to the previous choice exerted a significantly stronger influence on the drift rate than dissimilar samples (Fig. 2H). Notably, the weighting profiles in P2 and P3 deviated systematically from those in P1, for which no such confirmatory pattern was observed. The regression model also allowed the starting point (z) to vary as a function of the previous choice. Although the model captured an overall bias toward diagonal categorizations, previous choices did not reliably modulate the starting point (Fig. 2I). Collectively, these findings indicate that repetition biases primarily arise from asymmetric weighting during evidence accumulation rather than from a pre-decisional response bias. Participants exhibit a robust tendency to repeat previous choices within sequences, and this behavior is driven by confirmation-like distortions during the integration of sensory evidence

### Confidence is a driver of Confirmation Bias

After characterizing the computational mechanisms underlying choice repetition, we investigated the extent to which cognitive factors modulate this bias. Drawing on previous findings that suggest confidence acts as a gate for evidence integration (Balsdon et al., 2020; Braun et al., 2018; Rollwage et al., 2020), we hypothesized that higher levels of certainty would amplify the tendency to remain consistent with a prior judgment. Given that Experiment 1 did not include explicit confidence reports, we utilized a well-established proxy derived from response times (RT), based on evidence that faster responses typically denote higher decision confidence (Braun et al., 2018; Drugowitsch et al., 2012;Moreno-Bote, 2010; Sanders et al., 2016; Urai et al., 2017). We fitted a GLMM incorporating this RT-derived confidence proxy (RT_conf-1_) from the first presentation (P1) to predict choice behavior in the subsequent repetitions (P2 and P3; Table S1). Our analysis revealed that RT_conf-1_ significantly predicted choice consistency; specifically, participants were more likely to repeat their initial decision when their P1 response was faster (β = 0.54, SE = 0.090, Z = 6.07, p < 0.001). To assess the validity of this index and estimate participants’ metacognitive abilities, we conducted Experiment 2, in which participants provided explicit confidence ratings after each decision. Furthermore, we investigated whether the maintenance of this bias depends on exposure to identical sensory sequences across repetitions— reflecting a reliance on redundancy—or if it persists when observers are presented with novel, independent samples sharing an identical mean decision variable (repeated vs non repeated sequences; see methods section). This allowed us to determine whether the metacognitive “gating” of confirmation bias is a general principle of evidence integration or if it is specifically tied to the reprocessing of redundant information. We replicated the primary findings of Experiment 1 (see Fig. S5), confirming that participants’ responses were biased toward previous choices. This effect was consistent with a shift in the drift criterion rather than a change in the starting point (Fig. S6). Notably, choices involving novel stimulus sequences with an identical mean decision variable followed a similar confirmation bias pattern to those involving identical repetitions. While participants manifested a stronger self-consistency bias when the stimuli were identical (Fig. S7A; Table S2), both conditions were best explained by an asymmetric weighting of sensory evidence indicative of a confirmatory bias (Fig. S7B). These results suggest that the perceptual confirmation bias mechanism is not stimulus-specific but rather reflects a top-down integrative bias. Given that results across both conditions were qualitatively identical—including participants’ confidence reports (F(1, 36) = 0.02, p = 0.899, ηG^2^ < 0.001)—we collapsed the “repeat” and “non repeat” trials for the remaining metacognitive analyses.

We analysed the impact of confidence on participants’ decisions. We first confirmed that the confidence proxy, RT_conf-1_, was positively correlated with participants’ subjective confidence reports (r = 0.186, p < 0.001). We then fitted a GLMM including confidence from the n-1 presentation as a regressor to test its influence on subsequent decisions. As anticipated and in consistence with previous studies (Peters et al., 2017; Rollwage et al., 2020) and RT_conf-1_ analyses results and in Experiment 1, our results showed that confidence is a strong driver of self-consistency biases (Table S3). This was evidenced by a significant three-way interaction between initial response, confidence, and repetition (β = 0.677, SE = 0.095, z = 7.10, p < 0.001). This interaction suggests that probability of repeating a previous response in P2 is significantly modulated by the level of confidence held in that initial decision (Fig. 3A).

**Figure 3.**
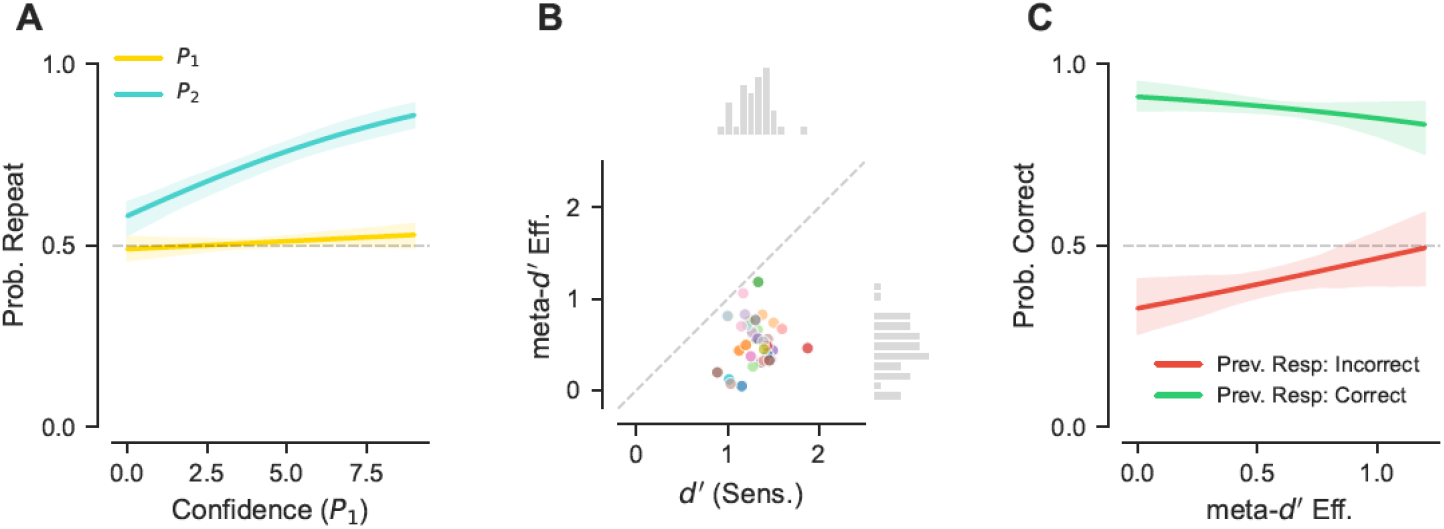
Metacognitive modulation of confirmation bias and adaptive decision revision. A, Probability of choice repetition as a function of subjective confidence in the preceding presentation. Higher subjective confidence in an initial decision significantly predicts an increased likelihood of repeating that choice in subsequent presentations, illustrating a confidence-gated self-consistency bias. B, Relationship between individual metacognitive efficiency (M_ratio_) and primary perceptual sensitivity (d’). Each colored data point represents a single participant, demonstrating that metacognitive efficiency is independent of individual perceptual sensitivity across the cohort. C, Probability of a correct final decision as a function of metacognitive efficiency, conditioned on initial performance in the first presentation (P1). The data are stratified by whether the participant was initially correct or initially incorrect in P1. While participants generally exhibit a strong tendency to repeat initially correct responses (green line), higher metacognitive efficiency significantly predicts the probability of recovering from an initial error (red line), highlighting the role of accurate metacognition in facilitating adaptive changes of mind. Confidence intervals (shaded regions) in panels a and c were calculated using a non-parametric bootstrapping approach with 500 permutations.

### Metacognitive efficiency ameliorates the impact of confirmation bias

We estimated each participant’s metacognitive efficiency (Fig. 3B) based on their task performance and confidence reports using a Hierarchical Bayesian modeling approach (Fleming, 2017; Fleming & Lau, 2014; Maniscalco & Lau, 2012). To test the influence of metacognition on decision-making, we extended our GLMMs to include metacognitive efficiency as a regressor for predicting participant choices (Table S4).

Crucially, we found a significant interaction between metacognitive efficiency and the previous response (β = −0.89, SE = 0.27, Z = −3.29, p < 0.001). This negative coefficient indicates that participants with higher metacognitive efficiency exhibited a significantly lower tendency to repeat previous responses. However, a critical question remains: is this reduction in repetition bias deployed strategically based on trial-by-trial performance, or is it a general trait that is insensitive to ongoing accuracy? To address this question, we modeled participants’ accuracy in each presentation as a function of their individual metacognitive efficiency and their performance (correct vs. incorrect) in the preceding presentation (Table S5). This analysis provides insight into how participants leverage their metacognitive abilities to improve performance and modulate confirmation bias in scenarios where information is repeated, specifically contrasting cases where they were initially correct versus those where they were wrong.

The model revealed a powerful two-way interaction between stimulus repetition and previous accuracy (β = 2.88, SE = 0.23, Z = 12.54, p < .001). This effect demonstrates a baseline confirmation bias: when participants were correct on their first encounter with a stimulus, repetition significantly boosted their accuracy on the second encounter. However, when they were initially incorrect, they showed a persistent tendency to repeat that error, leading to significantly lower accuracy upon repetition. This interaction effectively quantifies the “tripping twice over the same stone” phenomenon in our task.

Crucially, this baseline bias was modulated by a significant three-way interaction between stimulus repetition, metacognitive efficiency, and previous accuracy (β = −1.21, SE = 0.37, Z = −3.25, p = .001). This negative coefficient indicates that the impact of the confirmation bias is significantly reduced in participants with higher metacognitive efficiency (Fig. 3C).

While average performers struggle to self-correct after an initial error, those with high metacognitive efficiency appear to leverage their internal confidence signals to “break” the repetition cycle. This results in a strategic amelioration of the confirmation bias specifically when the previous choice was incorrect, allowing highly metacognitively efficient individuals to avoid repeating the same mistake.

## Discussion

To successfully interact with a dynamic world, the human brain must continuously sample sensory information to rapidly and flexibly update its beliefs about the current state of the environment. However, this imperative for flexibility clashes with a fundamental cognitive constraint: a pervasive bias toward self-consistency (Fischer & Whitney, 2014; Luu & Stocker, 2018; Urai et al., 2019). Perception is not a passive readout of the environment; it is actively sculpted by prior choices, creating a confirmation bias that permeates human cognition from high-level beliefs down to low-level sensory processing (Del Río et al., 2024; Kappes et al., 2019; Palminteri et al., 2017; Talluri et al., 2018). While prioritizing consistency may preserve cognitive resources in stable environments, it risks severe maladaptation in the highly dynamic, modern world (Kassin et al., 2013; Modgil et al., 2024; Zhou & Shen, 2022).

To investigate how the brain manages this trade-off, we developed a novel experimental paradigm that departs from traditional independent-trial decision-making designs. Instead, we provided participants with several repetitions of the exact same sensory information, allowing them to revise their previous choices. This approach aims to mimic the redundant sensory sampling characteristic of everyday environments. For example, when navigating heavy traffic, we frequently process the same perceptual features—such as the distance to a leading vehicle or the color of a traffic signal— repeatedly while shifting our attention across various sources. By utilizing a task in which information is highly temporally correlated, we naturally promote the emergence of confirmation biases, which can be computationally beneficial when an initial judgment is correct but detrimental when incorrect. This opens the following fundamental questions: How is information integrated across repeated experiences? Furthermore, what are the specific cognitive factors that determine our capacity to review and change previous choices when confronted with contradictory evidence? In this study, we successfully characterized the mechanisms by which observers integrate information across repetitions, providing empirical evidence that metacognitive skills are paramount in flexibly modulating the strength of this confirmation bias.

Our results confirmed that participants integrate information across successive presentations, as their performance improved with an increasing number of sequence repetitions. This effect manifested primarily as a strong bias toward their previous choices and was specifically tied to the awareness that information across repeated stimulus sequences was redundant; we found no significant evidence of similar biases across independent trial boundaries (e.g., from “P3” of one trial to “P1” of the next). More importantly, we demonstrated that observers accumulate a severe self-consistency bias. Computational modeling via Drift-Diffusion Models (DDMs) revealed this is not a superficial motor habit, but a profound distortion of sensory evidence integration. Participants actively overweighted confirmatory samples while exhibiting a selective, asymmetric blindness to contradictory evidence. We conceptualized this mechanism as an asymmetric lapse integrator, illustrating how the brain selectively drops inconsistent information to maintain a stable, albeit biased, perceptual narrative. Remarkably, this asymmetric weighting operates directly within the integration process itself. Within a single sequence, samples preceded by similar orientations exerted a stronger influence on the final decision (Fig. S4). Interestingly, this effect was not driven purely by physical similarity (e.g., raw orientation), but by similarity in the latent decision space (diagonality vs. cardinality). For example, physically orthogonal orientations that shared the same Decision Variable were weighted more heavily together. This establishes that the perceptual system transforms sensory inputs into a decision-relevant format prior to applying the consistency filter. Consequently, self-consistency is not a late-stage cognitive adjustment, but an automatic, deeply embedded computation that dictates perception across multiple temporal scales.

Systematically discarding contradictory evidence appears deeply maladaptive, posing a fundamental evolutionary paradox: how could such a rigid mechanism be adaptive? Our findings resolve this paradox by identifying metacognition as the critical regulatory system. We demonstrate that subjective confidence—whether implicit in reaction times or explicitly reported—acts as the gatekeeper for evidence accumulation. The integration of prior choices with new sensory data is computationally efficient only because confidence continuously modulates the strength of the confirmation bias (Balsdon & Philiastides, 2024; Braun et al., 2018; Peters et al., 2017; Rollwage et al., 2020). Critically, and consistent with previous simulation-based modeling work (Rollwage & Fleming, 2021), we provide empirical evidence that high metacognitive efficiency acts as a cognitive circuit breaker against irrational consistency. While most participants blindly compounded their errors by repeating initial choices, highly metacognitive individuals preserved their sensitivity to corrective evidence. When their initial choice was incorrect, these individuals successfully bypassed their own confirmation bias to revise their judgments upon re-exposure.

Our results demonstrate that confirmation bias is a fundamental feature of neural computation, balancing the need for stable perception with the computational cost of processing new data. However, this system relies on metacognitive skill to prevent adaptive consistency from degrading into rigid dogma. By identifying metacognitive efficiency as the critical cognitive circuit breaker against belief entrenchment, this work provides a mechanistic framework for understanding human susceptibility to misinformation. It suggests that in an increasingly redundant and polarized information landscape, societal resilience depends not on eliminating our inherent biases, but on cultivating the metacognitive capacity to regulate them. To fully realize this potential, future neuroimaging studies must now isolate the neural circuitry by which confidence signals dynamically gate confirmation bias, illuminating how the brain maintains flexibility in the face of redundant evidence.

## Methods

### Participants

For Experiment 1, 29 healthy adults (2 male, mean age = 20 years, SD = 2) with normal or corrected-to-normal vision were initially recruited. A distinct cohort of 39 healthy adults (6 male, mean age = 20 years, SD = 2.3) participated in Experiment 2. Two participants were excluded in Experiment 1 and one participant in Experiment 2, as their primary task sensitivity fell more than 2.5 standard deviations below the group mean. All participants provided informed consent prior to the commencement of the experiments, and all experimental protocols were approved by the local ethics committee. Participants received a baseline compensation of 10 euros for their time. To further incentivize optimal performance, participants in Experiment 2 could earn up to 2 additional euros via a lottery mechanism, wherein the probability of winning was dynamically linked to their objective accuracy in the primary decision-making task.

### Stimuli and Apparatus

Stimuli were presented on a 1920 × 1080 pixel monitor (53 cm wide) using PsychoPy (Peirce et al., 2019). Participants were seated at a viewing distance of 50 cm. The display refresh rate was 60 Hz (inter-frame interval = 16.67 ms). The stimuli consisted of sequences of six oriented Gabor gratings with a diameter of 14°, a spatial frequency of 2 cycles per degree, and a contrast of 0.5. Each sensory sequence was “sandwiched” between two masking events to prevent afterimage effects and clearly demarcate the stimulus stream. Each mask lasted 250 ms and was constructed from four overlapping gratings with orientations of 45°, 90°, and 135° at 0.5 opacity.

To ensure rigorous control over sensory evidence, a large matrix of stimulus sequences was pre-generated prior to the experiment. The orientation of each grating within a sequence was sampled from a von Mises circular distribution. These orientations were subsequently transformed into a Decision Variable (DV) using a sawtooth function defined as DV = sawtooth(4 · θ, 0.5), where θ represents the orientation in radians. This procedure mapped orientations to a range between −1, representing the cardinal axes (horizontal or vertical), and +1, representing the diagonal axes (± 45°). The generation process filtered out trials with average orientation standard deviations exceeding 1.5 units from the mean variability to ensure consistent internal sequence statistics. During the experiment, the software dynamically obtained the sequence for each trial by identifying all pre-generated sequences whose average DV fell within a narrow tolerance window (± 0.025) of the target difficulty level required by the adaptive staircase, and randomly selecting one sequence from this subset.

### Experimental Procedure

In both experiments, participants performed a two-alternative forced-choice (2AFC) task, judging whether the aggregate orientation of the stimulus sequence was closer to the cardinal or diagonal axes. To prevent motor habituation and decouple perceptual judgments from motor preparation, the spatial mapping of the response categories (i.e., whether the cardinal or diagonal icon appeared on the left or right side of the screen) was fully randomized for every individual presentation. Because the options constantly swapped, participants could not rely on habituated motor actions to repeat a choice, ensuring that any measured consistency bias reflected a true cognitive repetition rather than a motor carryover effect.

Experiment 1 was designed to establish individual perceptual thresholds and characterize the baseline consistency bias. Each trial began with a central fixation point presented for a randomized duration between 400 and 700 ms. A circular contour then appeared around the fixation point for 750 ms to signal the spatial location of the upcoming sequence. The identical physical stimulus sequence selected from the matrix was presented three consecutive times (P1, P2, and P3) per trial. Participants registered their perceptual decisions using a standard computer keyboard, pressing the “z” or “m” keys corresponding to the randomized on-screen spatial mapping (left and right respectively). Task difficulty was controlled using a 1-up/2-down adaptive staircase that adjusted the DV magnitude based exclusively on the accuracy of the P1 response to target an 80% accuracy threshold. Step sizes for the staircase were progressively reduced by a factor of 0.871 following correct responses to ensure stable convergence. Visual feedback for each presentation was provided via a 750 ms color change in the fixation point—green for correct and red for incorrect—only at the conclusion of the entire trial.

Experiment 2 focused on the modulatory role of metacognitive efficiency and restricted trials to two sequential presentations (P1 and P2). The initial fixation interval was extended to a range of 800 to 900 ms. In contrast to Experiment 1, participants in Experiment 2 registered all responses using a computer mouse. Following the stimulus presentation, participants made their perceptual categorization by clicking on a discrete, two-option visual slider, with the left/right configuration of the cardinal and diagonal options randomized per presentation. Immediately following each perceptual categorization, participants reported their subjective confidence using a second, continuous visual slider. Participants used the mouse to select a point along this scale, which was anchored by “Dudosa” (Doubtful) and “Segura” (Certain). To prevent spatial biases, the mouse cursor was automatically hidden and recentered to the middle of the screen at the start of each rating phase.

To isolate the effect of exact sensory redundancy on evidence integration, stimulus repetition was directly manipulated across four alternating blocks in Experiment 2. In “repeat” conditions, the exact physical sequence of six gratings selected for P1 was preserved and displayed again in P2. In the “non-repeat” control condition, the sequence shown in P2 was physically novel; the software randomly selected a new sequence from the pre-generated matrix that matched the exact same DV target value as P1. This ensured that the aggregate sensory evidence remained perfectly matched across presentations, but the local physical noise (the individual gratings) differed. To ensure high engagement with the confidence reporting, a lottery system was implemented where a random trial was selected at the end of each block, and participants were awarded points based on their objective accuracy in that specific trial.

### Computational Modeling and Data Analysis

To estimate the behavioral parameters, we fitted Generalized Linear Mixed Models (GLMMs) utilizing the pymer4 package in Python. Initially, we modeled the probability of participants categorizing a stimulus sequence as diagonal or cardinal using a logistic regression framework, specifying the sensory Decision Variable (DV) as the primary predictor. Subsequently, we incorporated the participant’s choice history by including the response from the immediately preceding presentation as an additional predictor. Notably, for the first presentation (P1) of a given trial, the preceding response was strictly defined as the final decision made in the prior trial. To computationally disentangle whether choice-history biases were driven by a pre-accumulation shift in the starting point or an asymmetric accumulation of evidence, we simulated reaction times and choices using synthetic drift-diffusion models. To simplify the modeling architecture, the synthetic experiment was designed to generate 5000 trials per iteration, with two consecutive presentations per trial. The simulated sensory stimulation consisted of a continuous sequence of evidence values scaled between −1 and 1, with an integrator accumulating this evidence stochastically over time. We simulated distinct integrators by applying a psychometric kernel to the evidence, dynamically adjusting its parameters to reflect three competing mechanistic hypotheses. First, to simulate a pre-accumulation response bias, the diffusion process was initialized with a starting point (z) shifted closer to the decision boundary of the previous choice. Second, to model an information-independent biased drift rate—representing a constant, uniform gain toward the previous response—we shifted the Point of Subjective Equality (PSE) of the observer’s psychometric function in the direction of the prior choice by modifying the detection threshold. Finally, to simulate an asymmetric integration mechanism wherein information consistent with a previous response is overweighted while inconsistent information is underweighted, we introduced a choice-dependent lapse rate to the psychometric function, thereby selectively dampening the accumulation of disconfirmatory evidence. After running a Monte Carlo simulation of 25 synthetic observers across these competing models, we qualitatively evaluated the resulting psychometric curves and reaction time (RT) distributions. We observed that only the asymmetric integration model (the lapse model) successfully reproduced the hallmark reduction in the slope of the choice-dependent psychometric functions—as well as the specific RT distribution patterns—that were observed in our empirical data during repeated stimulus exposures.

To empirically validate these mechanisms within our human cohort, we employed hierarchical Drift-Diffusion Modeling (HDDM) to jointly analyze choice accuracy and reaction times. The models parameterized the decision process through the drift rate (v), decision threshold (a), starting point ($z$), and non-decision time (Ter). Model parameters were estimated using Markov Chain Monte Carlo (MCMC) sampling with 2000 samples (burning the first 1000 samples). We rigorously inspected all HDDM model fits, MCMC traces, and posterior likelihood estimations to verify proper convergence and model specification. To adjudicate between competing models, we compared their Akaike Information Criterion (AIC) indices. Furthermore, to verify that individual samples within a sequence exerted a differential impact on the drift rate, we utilized a multiple linear regression HDDM framework, fitting the drift rate as a function of the DV distance of each sample relative to the previous choice. Specifically, we reordered the six sensory samples in each trial based on their similarity to the preceding decision, yielding six regressor columns. The first regressor column contained the samples most similar to the participant’s previous choice, while the final column contained the most dissimilar samples. To further investigate whether evidence integration was intrinsically modulated by the temporal continuity of the stimulus stream itself, we examined the influence of similarity between consecutive samples. We calculated the absolute difference in both orientation and DV between adjacent samples within each six-grating sequence, yielding five transition metrics per trial. Analogous to our previous analysis, these transitions were rank-ordered from most similar to most dissimilar relative to their immediate temporal predecessor. Finally, we incorporated these five ranked transition metrics as continuous regressors in a logistic GLMM to predict the participant’s final categorical decision (cardinal or diagonal), allowing us to systematically quantify the behavioral weight assigned to temporally continuous versus discontinuous sensory evidence.

Having established via DDM analyses that decision repetition biases were primarily driven by changes in the evidence integration rate, we reverted to the GLMM framework to simplify the modeling of metacognitive interactions. We fitted GLMMs incorporating our metacognitive variables as interaction terms with the repetition parameter to assess how metacognition modulates the consistency bias. We first derived an implicit measure of confidence from reaction times (RT_conf_). This approach consisted of log-transforming the reaction times, inverting the scale by subtracting each value from the maximum log-transformed RT computed within each participant and experiment, and subsequently z-scoring the results to yield a normalized, participant-level measure of decision certainty. To explicitly assess how confidence in a preceding decision influenced the subsequent choice, we lagged this metric by one presentation (RT_conf-1_) and incorporated it into our predictive models. In Experiment 2, we additionally utilized the explicit subjective confidence reports from participants as a regressor in the GLMM.

Finally, metacognitive efficiency was calculated utilizing participants’ primary task responses and explicit confidence reports. We employed subject-level Bayesian estimation using the metadPy toolbox in Python. This procedure yielded robust estimates of primary perceptual sensitivity (d’) and metacognitive sensitivity (meta-d). We subsequently derived the metacognitive efficiency metric (M_ratio_ = meta-d’/d’), which quantifies the degree to which an individual’s confidence ratings reliably track their objective accuracy while appropriately controlling for baseline differences in primary task performance. We modeled the probability of reporting cardinal or diagonal as a function of the DV, the previous choice, and its interaction with metacognitive efficiency. An additional logistic GLMM was utilized to model the probability of a correct response as a function of the correctness of the previous choice and individual metacognitive efficiency. To robustly calculate the confidence intervals for the model predictions, we employed a non-parametric bootstrapping approach with 500 permutations.

## Methods Ethics statement

The experiments were approved by the Bioethics committee (IRB00003099) of the University of Barcelona. All human subjects provided written informed consent.

## Acknowledgements

We thank Ángel Bujalance for his assistance during the initial stages of the project and Alexandre García-Durán and Jaime de la Rocha for their insightful discussions regarding the analyses. This work was supported by the Spanish Ministerio de Ciencia, Innovacio ń y Universidades, which is part of Agencia Estatal de Investigación (AEI), through the projects RTI2018-100977-J-I00 and RYC2022-037652-I to A.P.B., the projects PID2019-111199GB-I00 and PID2022-140426NB-I00 to L.F. (funded by MCIN/AEI/10.130 39/501100011033/ and FEDER a way to make Europe), and the Spanish Ministry of Science, Innovation and Universities (MICIU/AEI/10.13039/501100011033) and by “FEDER A way of making Europe” (ref: PID2023-146524NB), and by ICREA ACADÈMIA (2022) funded by the Catalan Institution for Research and Advanced Studies to R.M.B. The funders had no role in study design, data collection and analysis, decision to publish, or preparation of the manuscript.

