## Supplementary Figures for "Metacognitive Efficiency Reduces Confirmation Bias in Perceptual Decision Making"

**Simulation results:**

Below we report the results obtained from simulating synthetic data based on the different assumed integrators. We performed HDDM and model comparison on the synthetically simulated data to verify that the integrators behave as expected.

*Evidence Integrator with biased Starting point*

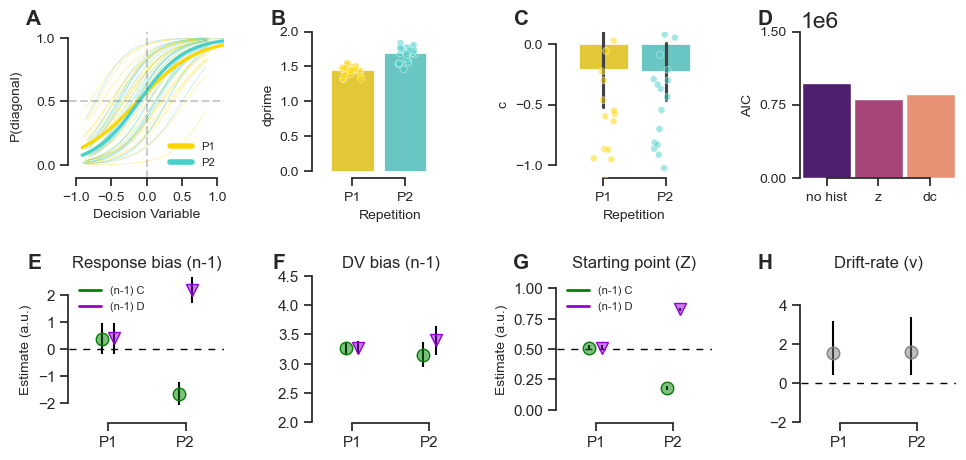

**Supplementary Figure 1 | Simulation of choice-history bias via starting point modulation.** A, Simulated psychometric curves for the first (P1, yellow) and second (P2, cyan) presentations. B–C, Signal detection theory (SDT) metrics (d' and c) derived from the simulated data. D, Model comparison (AIC) of HDDMs fitted to the simulated data. Note that for this simulation, the starting point (z) model provides the best fit. E, Response bias estimates (n-1) from the GLMM. F, Decision Variable (DV) bias estimates from the GLMM. G, Starting point (z) estimates from the HDDM, conditioned on the previous choice (green circle, cardinal; purple triangle, diagonal). This model successfully reproduces the horizontal shift in psychometric curves but fails to capture the empirical reduction in slope magnitude. H, Mean drift rate (v) across repetitions. Error bars represent 95% Bayesian credible intervals.

*Evidence Integrator with biased PSE*

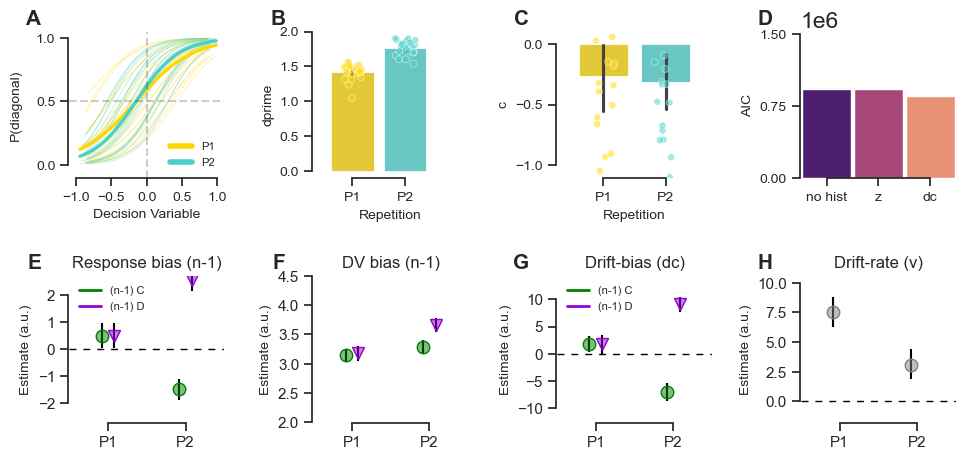

**Supplementary Figure 2 | Simulation of choice-history bias via Point of Subjective Equality (PSE) modulation**. A, Simulated psychometric curves for P1 (yellow) and P2 (cyan). B–C, Simulated SDT metrics (d' and c). D, Model comparison (AIC) indicating that the drift criterion (dc) model provides the superior fit for this simulation. E, Response bias estimates (n-1) from the GLMM. F, DV bias estimates from the GLMM. G, Drift bias (dc) estimates from the HDDM, conditioned on the previous choice. Modifying the PSE of the accumulator effectively shifts the psychometric function but, similar to the starting point model, does not reproduce the choice-conditioned loss in sensory precision observed in the human data. H, Mean drift rate (v) across repetitions. Error bars represent 95% Bayesian credible intervals.

*Evidence Integrator with asymmetric Lapses*

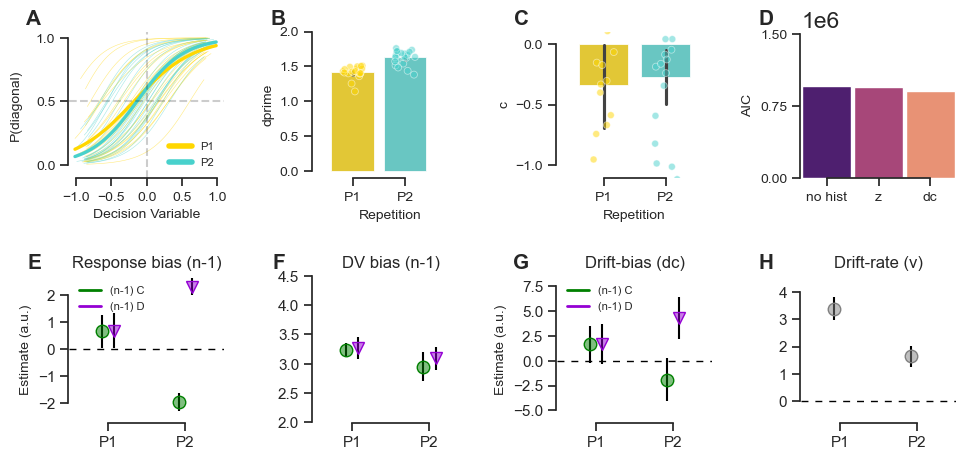

**Supplementary Figure 3 | Simulation of choice-history bias via asymmetric evidence integration (lapse rate).** A, Simulated psychometric curves for P1 (yellow) and P2 (cyan). B–C, Simulated SDT metrics (d' and c). D, Model comparison (AIC) adjudicating between the competing HDDMs. E, Response bias estimates (n-1) from the GLMM. F, DV bias estimates from the GLMM. G, Drift bias (dc) estimates from the HDDM. H, Mean drift rate (v) across repetitions. This simulation, which incorporates a choice-dependent lapse rate to dampen disconfirmatory evidence, is the only model that qualitatively reproduces both the horizontal psychometric shift and the hallmark reduction in the slope (sensory precision) observed in the empirical results. Error bars represent 95% Bayesian credible intervals.

**Experiment 1 stimuli kernels:**

We used a reverse correlation approach to estimate the temporal kernel of each sample within the sequence. We observed a recency effect, whereby the final samples exerted a stronger impact on the categorical decision. We also explored whether similarity (in terms of both DV and orientation) between consecutive sequence samples determined the contribution of each sample to the final choice. We found that during the integration process itself, DV similarity was determinant. Specifically, a sample preceded by a perfectly orthogonal oriented grating—which by definition belonged to the same category—had a stronger impact than a sample preceded by a partially dissimilar (45º) oriented stimulus. While we report the kernels for Experiment 1 here (Fig. S4), we replicated this finding in Experiment 2.

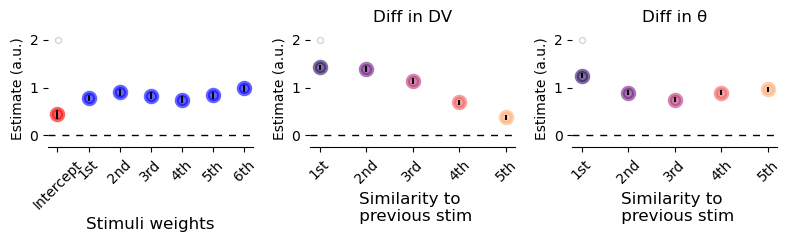

**Supplementary Figure 4 | Psychophysical kernels and temporal integration weights in Experiment 1.** Left, Estimated stimulus weights for each of the six gratings within a sequence, ordered chronologically. The intercept (red) represents the baseline bias, while blue circles indicate the regression weights for each temporal position, illustrating a relatively stable contribution of each sample to the final decision. Middle, Influence of sensory continuity on choice, quantified by the similarity in Decision Variable (DV) between consecutive samples. Weights are ranked from most similar (1st) to least similar (5th) to the preceding stimulus. The monotonic decrease in estimate magnitude suggests that information more consistent with the immediate sensory past exerts a stronger influence on the current categorization. Right, Influence of sensory continuity on choice quantified by the angular difference (theta) between consecutive samples. Transition weights are ranked by similarity (1st to 5th). Consistent with the DV analysis, gratings that are more similar in orientation to the previous sample are assigned higher weights during evidence integration. Error bars represent 95% confidence intervals.

**Experiment 2 results:**

In Experiment 2, we replicated the primary results observed in Experiment 1. Participants' sensitivity increased significantly with the number of repetitions. We also found that psychometric curves were strongly biased toward previous choices. As in Experiment 1, the bias in P2 was associated with a reduction in the psychometric curve sensitivity conditioned on previous responses (Fig. S5H). This pattern is better explained by an integrator with lapses (Fig. S3F) than by a biased integrator (Fig. S2F).

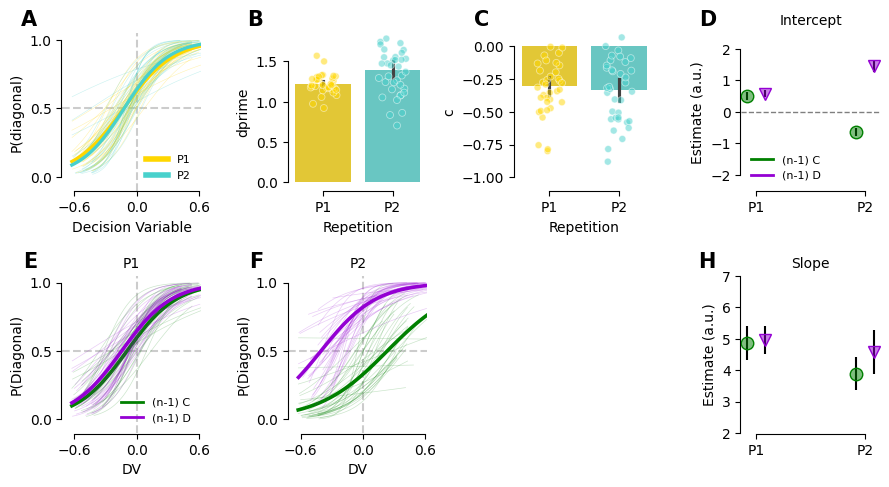

**Supplementary Figure 5 | Behavioral performance and choice-conditioned psychometric shifts for Experiment 2.** A, Psychometric curves illustrating the probability of a "diagonal" categorization as a function of the Decision Variable (DV) for the first (P1, yellow) and second (P2, cyan) presentations. Thin lines represent individual participants, while thick lines indicate the group-level logistic fit. B–C, Signal detection theory (SDT) metrics across repetitions. Sensitivity (d'; B) increased between presentations, whereas the decision criterion (c; C) demonstrated a consistent negative bias toward the cardinal category. Points represent individual participant data; bar heights represent the mean, and error bars indicate s.e.m. D, Intercept estimates from the hierarchical mixed-effects model, quantifying systematic bias as a function of choice history. Estimates are conditioned on whether the previous response was cardinal (green circles) or diagonal (purple triangles), with error bars representing 95% confidence intervals. The significant divergence at P2 indicates that the preceding choice exerts a robust influence on the baseline response probability. E–F, Choice-conditioned psychometric curves for the first (E) and second (F) presentations. Data are stratified by the observer’s choice in the immediately preceding presentation (n-1). Note the substantial horizontal shift at P2 between cardinal-conditioned (green) and diagonal-conditioned (purple) curves, capturing the impact of the previous judgment on current perception. H, Slope estimates from the hierarchical mixed-effects model, representing sensitivity to the sensory DV across repetitions, conditioned on the previous response. Similar to Experiment 1, a reduction in slope magnitude is observed at P2, indicating a loss in precision when sensory information is repeated.

In Experiment 2, we replicated the primary drift-diffusion modeling results observed in Experiment 1. Decision biases were better explained by a biased drift criterion (Fig. S6A). Furthermore, sensory samples that were more consistent with the participant's previous choice exerted a stronger impact on the drift rate compared to those that were less consistent Fig. S6D).

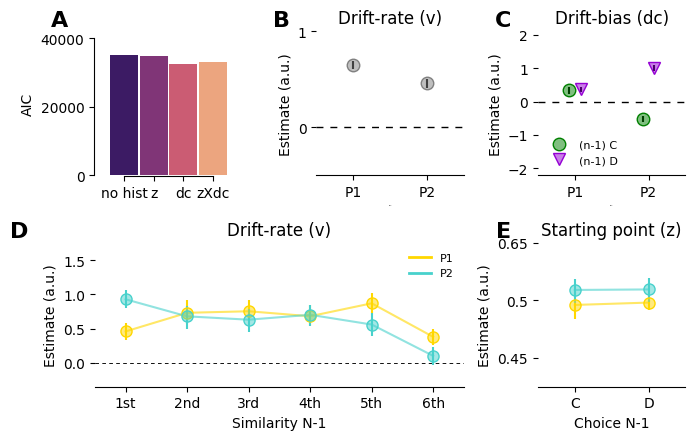

**Supplementary Figure 6 | Computational mechanisms of choice-history bias during repeated stimulus exposure in Experiment 2.** A, Model comparison using the Akaike Information Criterion (AIC). Hierarchical Drift-Diffusion Models (HDDMs) were compared to adjudicate between distinct mechanisms of history integration. Similar to Experiment 1, the model incorporating history-dependent shifts in the drift criterion (dc) provided the best fit, outperforming models assuming no history dependence ("no hist"), a history-dependent starting point (z), or a combination of both (zXdc). B, Estimated mean drift rate (v) across the first and second presentations (P1, P2). Accumulation speed decreased upon repeated exposure to the stimulus. C, Drift bias (dc) estimates conditioned on the observer’s previous choice (n-1): cardinal (C, green circles) or diagonal (D, purple triangles). The accumulation process became increasingly biased toward the boundary of the preceding choice during the second presentation (P2). D, Evolution of the drift rate (v) as a function of sensory sample similarity to the previous choice, ranked from most similar (1st) to least similar (6th). For both P1 (yellow) and P2 (cyan), sensory evidence consistent with the previous judgment was assigned higher accumulation weights. E, Starting point (z) estimates conditioned on the previous choice. Across both P1 and P2, the starting point remained near the unbiased baseline (0.5), though it was significantly closer to the diagonal boundary in the P2 condition. Nevertheless, previous choices did not affect the starting point, indicating that choice history modulates the rate of evidence integration rather than pre-accumulation offsets.

In Experiment 2, we also investigated whether repeated sequences had to be identical to produce a confirmation bias. In different blocks, participants experienced either the exact same sequence repeated or a different sequence with the same mean Decision Variable (DV). Participants were aware of which block they were in and what the manipulation entailed. Although participants exhibited a tendency to repeat their previous choice in both block types, we found they were more biased toward repeating the same response in P2 when the sequence was identical. Nevertheless, using a linear regression HDDM, we observed that the drift rate was biased according to a confirmation bias pattern in both non-repeated and repeated sequences, although the effect was significantly stronger in the repeated condition. Whether the increase in repetition bias in repeated compared to non-repeated sequences is due to low sensory redundancy, or because participants had prior knowledge that the stimuli sequence was being exactly repeated, cannot be answered in the current study.

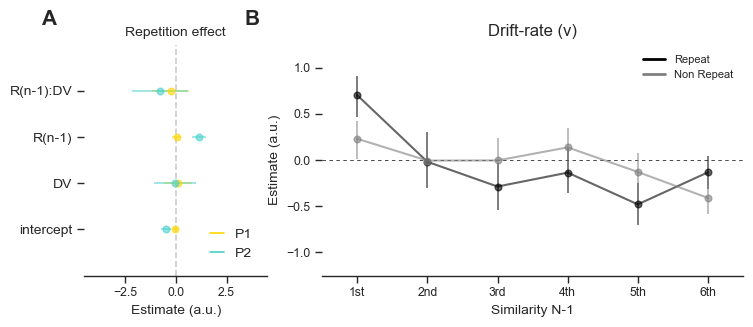

**Supplementary Figure 7 | Repetition effects on evidence integration parameters in Experiment 2.**A, Repetition effect estimates for GLMM parameters contrasted against the first presentation (P1). The plot illustrates the interaction of stimulus repetition with the change in the intercept, average Decision Variable (DV), preceding response (R(n-1)), and their interaction (R(n-1):DV) during the second presentation (P2, cyan) relative to P1 (yellow). Error bars represent 95% confidence intervals. These results suggest a significantly stronger tendency to repeat a decision when the sequences are identical. B, Evolution of the drift rate (v) as a function of sensory sample similarity to the previous choice, ranked from most similar (1st) to least similar (6th). Estimates are shown for the second presentation of trials where the previous choice was repeated (black line) versus non-repeated (gray line). The presented values are relative to the estimated P1 drift rates. The divergence at the first and sixth samples suggests that the weighting of sensory evidence is modulated by choice consistency in P2. This same pattern appears in both repeat and non-repeat conditions, indicating that confirmation bias is ubiquitous across repetition conditions; however, the magnitude of the bias is stronger in the exact repetition condition.

**GLMM Tables**

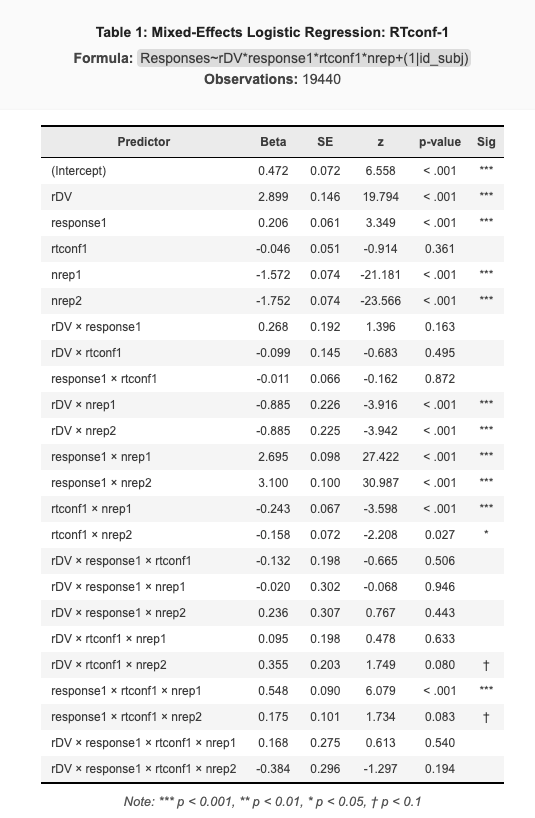

**Supplementary Table 1 | Mixed-effects logistic regression analysis of decision certainty in Experiment 1.** The table presents the fixed-effects estimates from a generalized linear mixed model (GLMM) predicting participant responses (cardinal or diagonal). The model includes the sensory Decision Variable (rDV), the preceding choice (response1), and an implicit measure of confidence in derived from reaction times in the previous presentation (rtconf1) as primary predictors. To assess how these influences evolve over time, interaction terms were included with the repetition number (nrep1, nrep2). Beta coefficients, standard errors (SE), z-values, and p-values are reported for all main effects and interactions. Random intercepts were included for each participant (id\_subj) to account for individual variability. Statistical significance is denoted as follows: *** p < 0.001, p < 0.01, * p < 0.05, † p < 0.1.

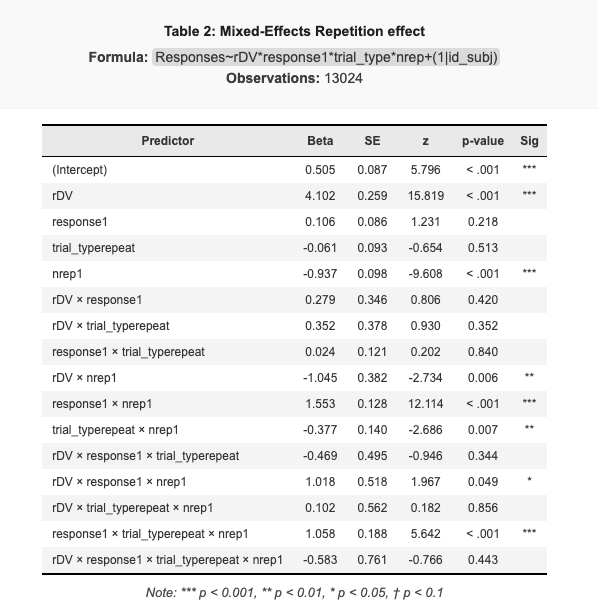

**Supplementary Table 2 | Mixed-effects logistic regression: Interaction between trial type (non repeated/ repeated) and choice repetition.** The table presents the fixed-effects estimates from a generalized linear mixed model (GLMM) predicting participant responses. The model evaluates the influence of the sensory Decision Variable (rDV), the preceding choice (response1), trial type (trial_type), and the repetition index (nrep1). Interaction terms were included to assess how the repetition of identical stimulus sequences (repeat trials) modulates the choice-history bias compared to non-repeated sequences. Beta coefficients, standard errors (SE), z-values, and p-values are reported for all main effects and interactions. The model includes random intercepts for each participant 1 | id_subj) to account for individual variability. Statistical significance is denoted as follows: *** p < 0.001, p < 0.01, * p < 0.05, † p < 0.1.

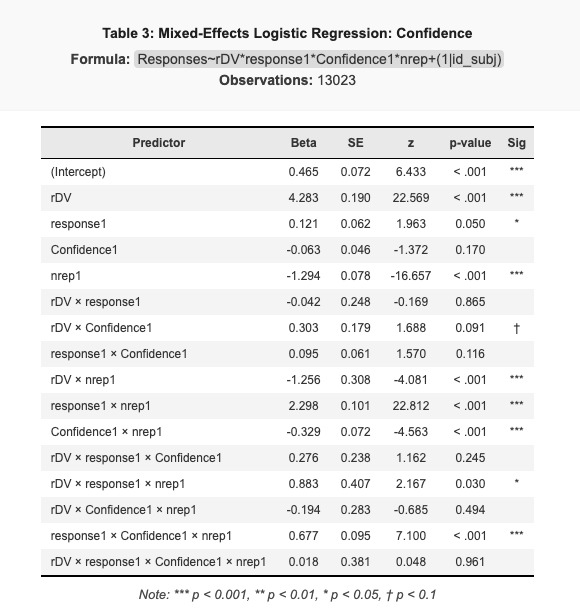

**Supplementary Table 3 | Mixed-effects logistic regression analysis of subjective confidence and choice repetition in Experiment 2.** The table presents the fixed-effects estimates from a generalized linear mixed model (GLMM) predicting participant responses. The model evaluates the influence of the sensory Decision Variable (rDV), the preceding choice (response1), and explicit subjective confidence reports from the previous presentation (Confidence1). Interaction terms were included with the repetition index (nrep1) to assess the temporal evolution of these effects. Beta coefficients, standard errors (SE), z-values, and p-values are reported for all main effects and interactions. The model includes random intercepts for each participant (id\_subj) to account for individual variability. Statistical significance is denoted as follows: *** p < 0.001, p < 0.01, * p < 0.05, † p < 0.1.

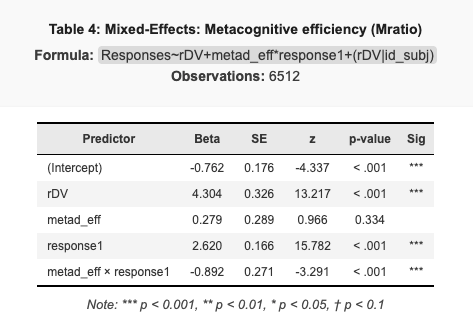

**Supplementary Table 4 | Mixed-effects logistic regression: Interaction between metacognitive efficiency and choice history in Experiment 2.** The table summarizes the fixed-effects estimates from a generalized linear mixed model (GLMM) predicting categorical responses (cardinal or diagonal). The model examines the influence of the sensory Decision Variable (rDV), individual metacognitive efficiency (metad_eff), and the preceding choice (response1). A key interaction term (metad_eff * response1) was included to assess how metacognitive ability modulates the impact of choice history on current decision-making. Beta coefficients, standard errors (SE), z-values, and p-values are provided for all predictors. The model incorporates random intercepts and slopes for the Decision Variable grouped by participant (rDV | id_subj) to account for hierarchical data structures. Statistical significance is indicated as follows: *** p < 0.001, p < 0.01, * p < 0.05, † p < 0.1.

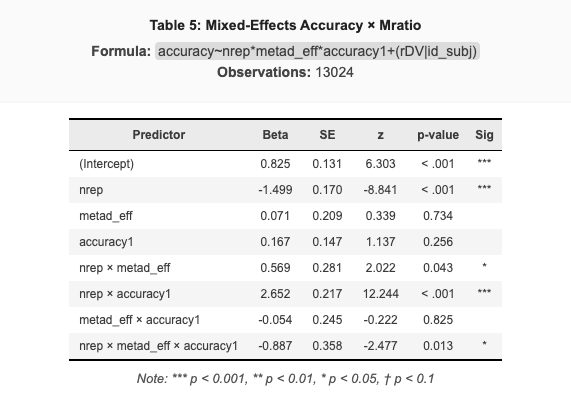

**Supplementary Table 5 | Mixed-effects logistic regression: Interaction between performance, stimulus repetition, and metacognitive efficiency in Experiment 2.** The table summarizes the fixed-effects estimates from a generalized linear mixed model (GLMM) predicting decision accuracy (correct or incorrect). The model evaluates the relationship between stimulus repetition, individual metacognitive efficiency (metad_eff), and initial accuracy in the first presentation (accuracy1). Interaction terms were included to assess how metacognitive efficiency modulates the probability of maintaining correct responses or recovering from initial errors across repetitions. Beta coefficients, standard errors (SE), z-values, and p-values are provided for all predictors. The model incorporates random intercepts and slopes for the Decision Variable grouped by participant (rDV | id_sub) to account for individual variability in sensory sensitivity. Statistical significance is indicated as follows: *** p < 0.001, p < 0.01, * p < 0.05, † p < 0.1.
